# Asymmetry of the temporal code for space by hippocampal place cells

**DOI:** 10.1101/060590

**Authors:** Bryan C. Souza, Adriano B. L. Tort

## Abstract

Hippocampal place cells convey spatial information through spike frequency (“rate coding”) and spike timing relative to the theta phase (“temporal coding”). Whether rate and temporal coding are due to independent or related mechanisms has been the subject of wide debate. Here we show that the spike timing of place cells couples to theta phase before major increases in firing rate, anticipating the animal’s entrance into the classical, rate-based place field. In contrast, spikes rapidly decouple from theta as the animal leaves the place field and firing rate decreases. Therefore, temporal coding has strong asymmetry around the place field center. We further show that the dynamics of temporal coding along space evolves in three stages: phase coupling, phase precession and phase decoupling. These results suggest that place cells represent more future than past locations through their spike timing and that independent mechanisms govern rate and temporal coding.

## INTRODUCTION

The rodent hippocampus plays a role in spatial memory and navigation (Buzsáki and Moser, 2013; Morris et al., 1982). Some hippocampal neurons, called place cells, increase their firing rate when the animal is at a specific location of the environment, known as the ‘place field’ of the cell (O’Keefe and Dostrovsky, 1971). As the animal crosses place fields, place cells form spike sequences coordinated by the hippocampal theta rhythm (~5-12 Hz) by firing action potentials coupled to earlier phases of the cycle, a phenomenon known as ‘phase precession’ (O’Keefe and Recce, 1993). Place fields and phase precession are considered canonical examples of rate and temporal coding, respectively, in which the firing rate of the neuron and the exact spike timing relative to the theta cycle provide information about space (Jensen and Lisman, 2000; O’Keefe and Burgess, 2005; Wilson and McNaughton, 1993). Whether temporal and rate coding are governed by independent or related mechanisms has been widely debated (Cei et al., 2014; Harris et al., 2002; Huxter et al., 2003; Mehta et al., 2002). For instance, Harris et al. (2002) showed that changes in firing rate predict changes in spiking theta phase, and concluded that temporal and rate coding are intrinsically related. On the other hand, Huxter et al. (2003) showed that the theta phase of spiking correlates more with animal position than with firing rate, and suggested that place cells use two independent spatial codes. More recently, by simultaneously controlling for firing rate and position within the place field, Cei et al. (2014) demonstrated that spiking phase depends on both firing rate (for a fixed position) and position (for a fixed firing rate), thus seemingly conciliating the discrepant conclusions in Harris et al. (2002) and Huxter et al. (2003). However, here we report a key new piece of evidence showing that temporal and rate coding dissociate to a much greater extent than previously recognized in these studies.

Since temporal coding requires the coupling of place cell spikes to theta phase (Harris et al., 2002; Huxter et al., 2003), we revisited the rate vs temporal coding debate by investigating the spatial evolution of theta coupling strength. Surprisingly, we found that the spike timing of place cells couples to theta before major increases in firing rate, that is, before the animal enters the classical, rate-based place field. In contrast, theta-phase coupling rapidly ceases as the animal leaves the place field and firing rate decreases. These results reveal that temporal coding has strong asymmetry around the place field center; moreover, since the spatial receptive field of temporal coding is larger before than after the place field center, they further suggest that place cell spike timing may code more for upcoming than past positions. Interestingly, we also found that place cells do not couple to theta phase at positions distant from the place field center, and that the dynamics of temporal coding along space separates into three stages: phase coupling, phase precession and phase decoupling. These findings shed new light on how place cells represent space, and suggest independent mechanisms of temporal and rate coding.

## RESULTS

### Rescaling space by place field length

We analyzed 1071 place fields from 689 place cells recorded from the hippocampal CA1 region of three rats while they ran back and forth on a linear track. Figure 1A shows spike locations for a place cell along with the animal’s trajectory (top) and the cell’s place field (bottom). As in previous work (Harris et al., 2002; Senior et al., 2008), we estimated place field length from a firing rate threshold, while the position of maximum firing defined the place field center. Figure 1B shows raw and theta-filtered local field potentials (LFPs) together with spike times during a place field crossing; notice the characteristic precession of spike timing within theta cycles (white/gray shades) from ascending to descending phases. Phase precession is most evident when plotting the theta phase of spiking as a function of position for multiple runs (Figure 1C, top). To study the relation between spiking phase and space in a systematic way, we first normalized place fields to account for their variable lengths and locations. To that end, the position of each spike was expressed as the animal’s distance to the place field center divided by the place field length (Figure 1C, middle). This procedure allowed visualization of the theta phase of spiking as a function of relative distance in units of place field length, centered at the peak activity of the place cell. Finally, as in previous work (Cei et al., 2014; Mehta et al., 2002) we binned theta phase and space and expressed spike counts by means of heat maps (Figure 1C, bottom).

**Figure 1.**
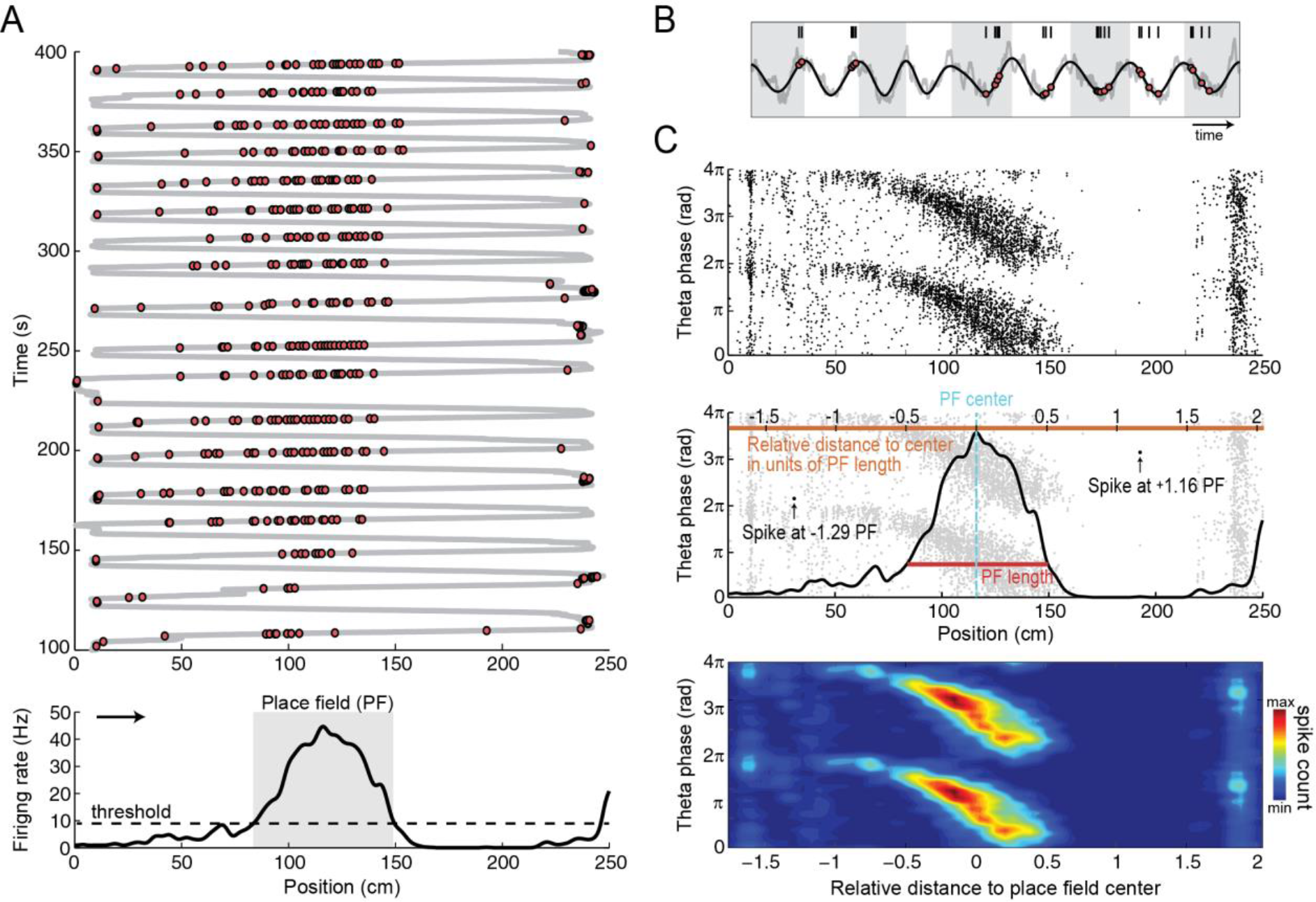
Spatial normalization by place field length. (A) Top, trajectory of a rat running back and forth on a linear track (gray line); red circles represent spikes of a place cell. Bottom, firing rate of the same cell as a function of position during runs to the right (arrow). The dashed line denotes the threshold used to estimate the length of the place field (gray shade). (B) Example of phase precession during a run to the right. Gray/white shades mark theta cycles. (C) Top, black dots represent all spikes of the cell in A as a function of position in cm (x-axis) and theta phase (y-axis). Middle, scheme of spatial normalization. Light gray dots represent spikes. The center of the place field was defined as the position of maximum firing rate. The orange horizontal scale on top shows the distance to the place field center in units of place field length. Arrows highlight the relative distance of two spikes. Bottom, heat map representation of spike counts per relative distance and theta phase.

### Theta coupling of place cells preceding place field entrance

Figure 2A,B shows the mean spike count per bin of theta phase and relative distance to the place field center (n= 1071 place fields). In these maps, the phase distribution of place cell spiking cannot be properly visualized at space bins distant from the place field center because of their much lower spike count (Figure 2C). To circumvent this, we normalized spike counts within each space bin (a column slice) by its mean over all phases (Figure 2D). This space-normalization allowed for assessing the phase distribution of spikes at positions away from the place field center. Surprisingly, we found that spikes concentrate near the theta peak before the animal enters the place field, at positions where place cells exhibit no major changes in firing rate (compare Figure 2D with 2B,C). Consistently, Figure 2E shows a sharp increase in spike-phase coupling strength preceding a slower increase in firing rate; spike-phase coupling transiently decreases as firing rate peaks due to bursting at the place field center, and later returns to basal levels as firing rate decreases. The fact that spike timing strongly couples to theta phase before changes in firing rate suggests dissociated mechanisms of rate and temporal coding.

**Figure 2.**
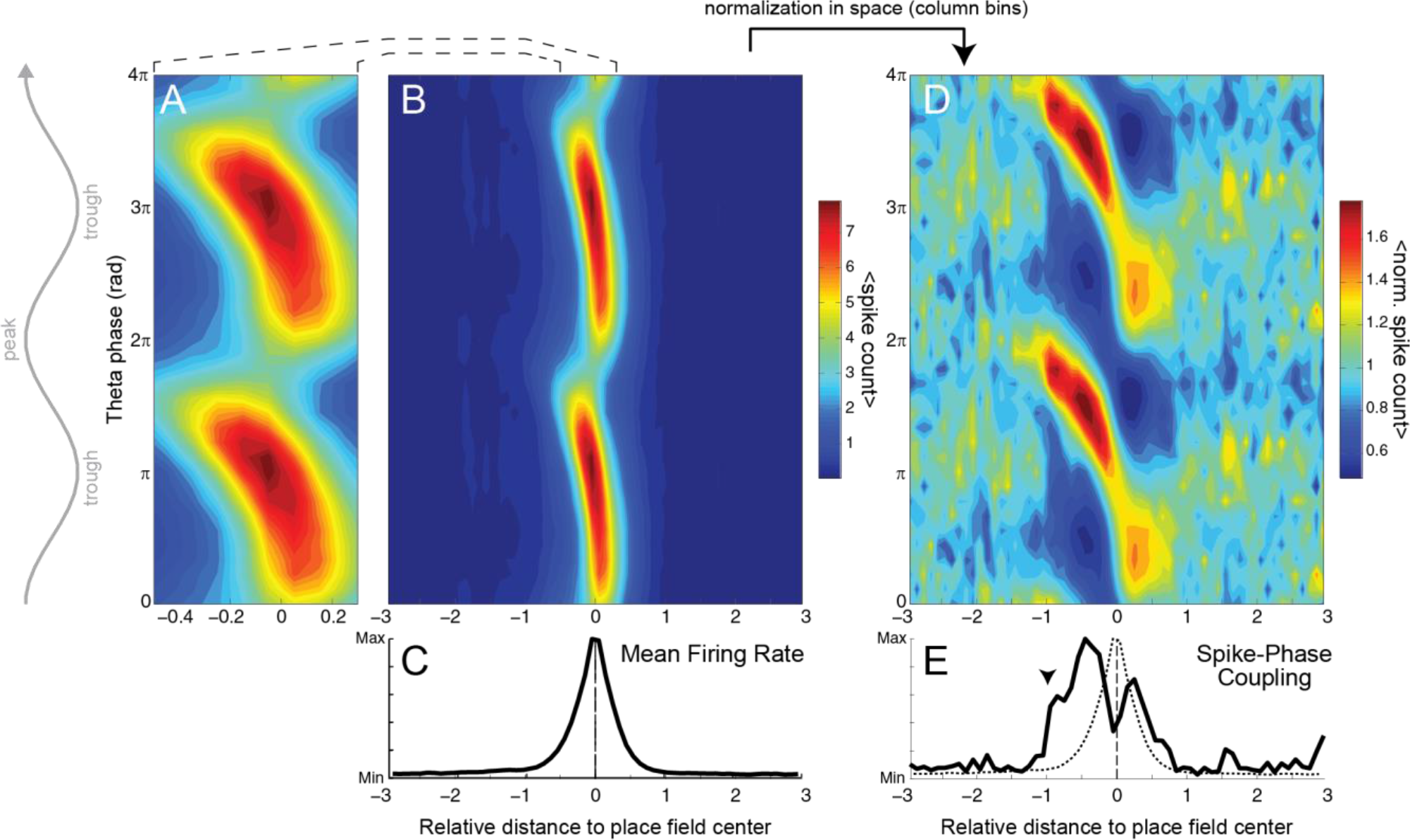
Place cells couple to theta phase before increasing their firing rate. (A) Average spike count per relative distance and theta phase (n=1071 place fields), showing the characteristic phase precession (2*π* is the peak of theta recorded from the CA1 pyramidal cell layer). (B) Zoomed-out view of the data in A. (C) Mean firing rate as a function of relative distance. (D) Same data as in B, normalized by the mean number of spikes at each position. (E) Spike-phase coupling strength per relative distance (solid line); the dashed line reproduces the firing rate in C. Notice that place cell spikes align to a preferred theta phase before major increases in firing rate (arrowhead).

To assess whether the results could be due to differences in spike counts among place fields, we next calculated the mean spiking phase for each place field per bin of relative distance. Figure 3A shows the spatial histogram of mean spiking phases. In each histogram column (space bin), individual place fields contribute with a single count at the place cell’s mean spiking phase, irrespective of the number of spikes. Columns with nonuniform phase distributions indicate that place cells have similar mean spiking phase at these positions, with warmer colors denoting preferred theta phases. We next computed theta-phase coupling strength (TPC) using the distribution of mean spiking phases, and used a surrogate procedure to delimit statistically significant values (black arrows in Figure 3A). TPC was above chance much before changes in mean normalized firing rate (FR) (Figure 3B) and followed a similar time-course as the theta-phase coupling of pooled spikes (Figure 2E), extending our findings from the spike to the place cell level. In fact, such a phenomenon of strong theta-phase coupling before a major increase in firing rate is also apparent in individual cells; for instance, notice that the example place cell in Figure 1C aligns its spikes to the theta peak at ~50 cm, much before its peak firing rate at ~115 cm (see Figure S1 for other examples). This result also holds true when analyzing each of the three rats separately (Figure S2). Interestingly, and in contrast to previous views (Bose et al., 2000), Figures 2E and 3B also demonstrate that the coupling of place cells to theta phase is spatially transient, which is to say that TPC – as FR – has a spatial receptive field. Moreover, they further show that different coupling strengths may be associated to the same firing rate depending on the animal’s position (e.g., compare FR and TPC at the relative distance of −1 and +1).

**Figure 3.**
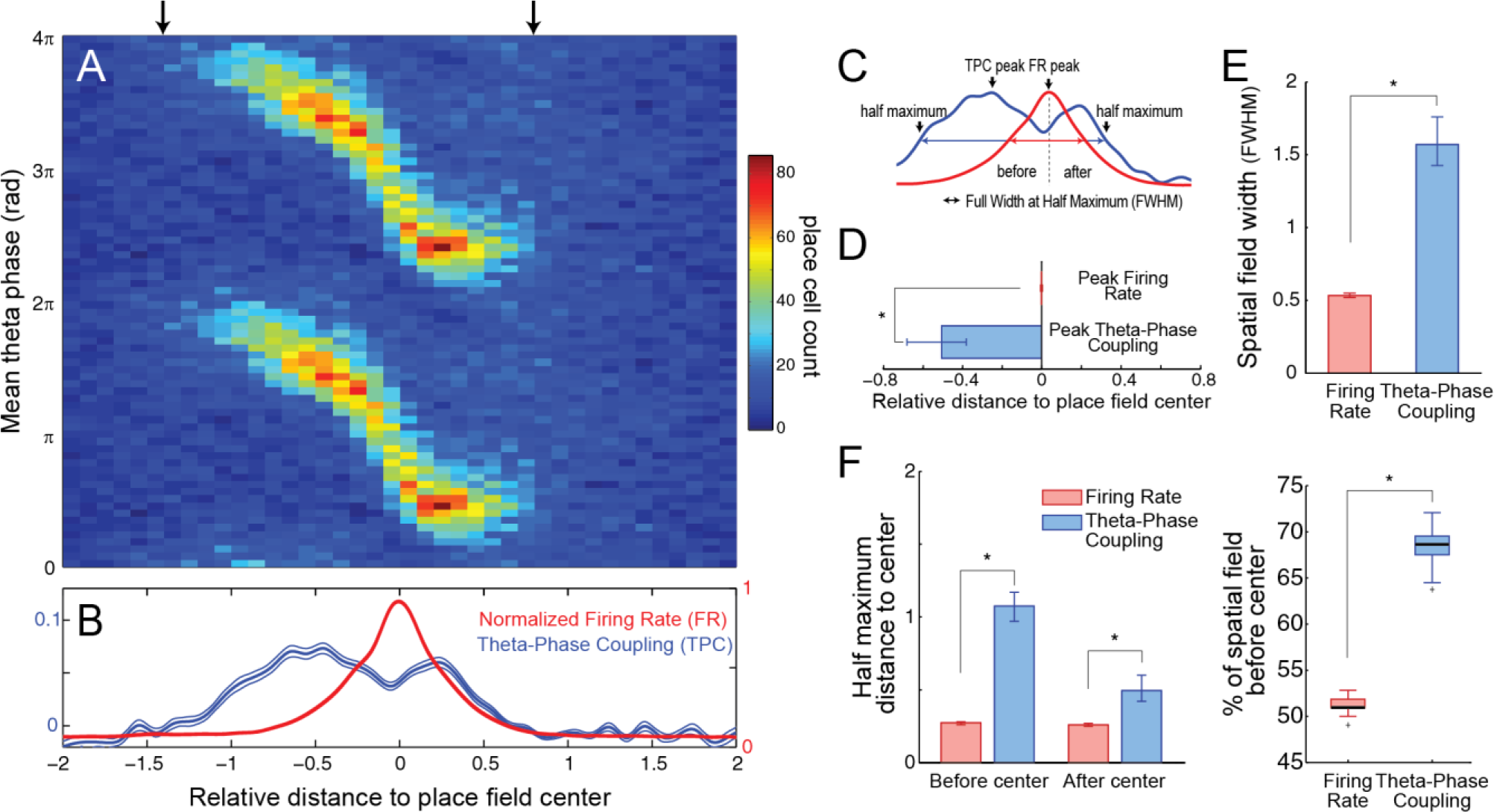
Asymmetry of the temporal code for space by hippocampal place cells. (A) Histogram of mean spiking phase. At each position, individual place fields contribute with one count to the phase bin of their place cell’s mean spiking. Arrows mark the region where the distribution of mean spiking phases statistically differs from the uniform distribution, indicating coordinated theta-phase coupling across cells. (B) Theta-phase coupling strength (TPC, blue) and mean normalized firing rate (FR, red) as functions of space. Thin lines denote 99% confidence intervals. The TPC curve was corrected by subtracting chance values (see Material and Methods) (C) Scheme describing the identification of peak and half maximum values as well as of the full width at half maximum (FWHM) for TPC and FR curves. (D) Mean relative distance of TPC and FR peaks, showing that place cells are maximally modulated by theta phase before their peak firing rate. (E) Comparison of spatial field width when defined by the FWHM of FR vs TPC. (F) Relative distances of FR and TPC half maximum values to the place field center (left) and percentage of spatial field located before the center when defined by FR vs TPC FWHM (right). In D-F, the error bars denote 99% confidence interval of the mean; *p<0.01.

### Asymmetry of the temporal code for space

To further characterize differences in temporal and rate coding, we computed the peak and the full width at half maximum (FWHM) of the TPC and FR curves along space (Figure 3C). TPC peaked as rats entered the place field (mean peak distance: −0.51; CI_99_: [−0.68;−0.38]), much before the FR peak at the place field center (Figure 3D). The FWHM of the TPC curve was significantly wider than that of the FR curve, implying that spatial fields would be larger if defined by changes in theta-phase coupling (mean FWHM for FR: 0.53; CI_99_: [0.52;0.55]; for TPC: 1.56; CI_99_: [1.42;1.79]; Figure 3E). Moreover, while the portion of the FR curve above its half-maximum value was roughly symmetrical around the place field center (see also Figures S3 and S4), most of TPC FWHM occurred before the center (Figure 3F; % FWHM before center for FR: 51.4; CI_99_: [50.0; 52.8]; for TPC: 68.5; CI_99_: [65.1;71.6]). These results show that hippocampal place cells have an asymmetric temporal code for space.

It has been previously reported that rate-based place fields become skewed with experience: in familiar tracks, place cells spike more at locations preceding the location of their peak firing rate (Mehta et al., 2000; Lee et al., 2004). Such an effect is better noticed when place field width is defined at 10% of the peak firing rate; on the other hand, place field skewness is not significantly different from 0 if computed at 50% of the peak firing rate (see Figure 2 in Mehta et al., 2000). Consistent with this, here we found that at 50% of the peak firing rate (i.e., at the FWHM), rate-based place fields were roughly symmetrical (Figures 3, S3 and S4), while FR skewness was significantly different from zero at 10% of the peak (−0.139 ± 0.018, p<0.001; Figure S3) and similar in magnitude to values reported in Lee et al. (2004) for CA1 place cells (−0.09 ± 0.04 to −0.12 ± 0.04, their Table 2). Importantly, however, the percentage of the spatial receptive field before the place field center was much larger for TPC than FR at all thresholds from the peak value (Figure S4), which is to say that temporal coding has much greater asymmetry around the place field center than rate coding even at thresholds in which FR is negatively skewed.

Interestingly, the joint dynamics of FR and TPC revealed that space coding by place cells occurs in three stages, and that phase precession is an intermediate stage in their temporal organization (Figure 4). In the first stage, place cells align to a fixed theta phase while maintaining a low spiking rate; this ‘phase coupling stage’ takes place before the animal enters the place field. Next, during the ‘phase precession stage’, place cells spike at higher rates and at sharply varying phases of the theta cycle as the animal traverses the place field. Finally, in the third stage, place cells decouple from theta while spiking at decreasing rates at the same phase reached after precessing; this ‘phase decoupling stage’ occurs while the animal leaves the place field.

**Figure 4.**
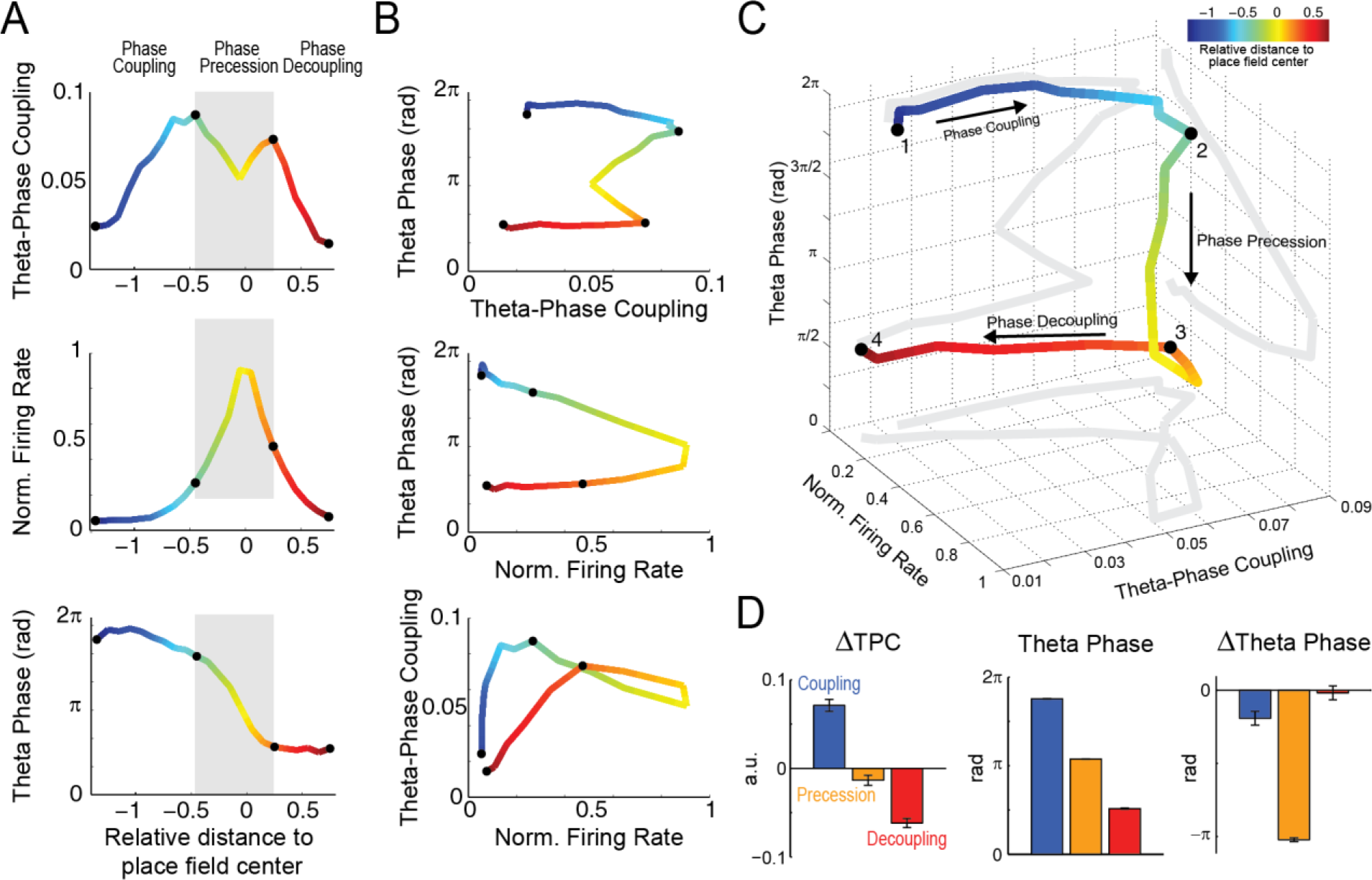
Three-stage model of space coding by place cells. (A) Theta-phase coupling strength (top), mean normalized firing rate (middle) and mean theta phase of spiking (bottom) per relative distance. Gray shaded area separates place cell activity into three stages, as labeled. (B) 2D plots of all pairwise combinations of the independent variables in A. (C) 3D plot of theta-phase coupling strength, firing rate and mean spiking theta phase. Gray lines show 2D projections (same curves as in B). Place cells align to a preferred theta phase before major changes in firing rate and spiking phase (“Phase Coupling”; 1 – 2). Next, the spiking phase rapidly advances as firing rate peaks (“Phase Precession”; 2 – 3). Finally, theta-phase coupling and firing rate return to basal levels while spiking occurs at the same theta phase reached after the precession (“Phase Decoupling”; 3 – 4). (D) Mean change in theta-phase coupling strength (left), mean theta phase of spiking (middle), and mean change in spiking phase (right) during each stage. In A-C, the line color denotes the relative distance to the place field center and black dots mark stage boundaries.

## DISCUSSION

Hippocampal place cells convey spatial information through both firing rate and spike timing. In the first case, firing rate increases as the animal approaches a specific location in space known as the place field of the cell (O’Keefe and Dostrovsky, 1971) (Figure 1A). In the second, place cells spike at varying phases of the field theta oscillation (~5-12 Hz), gradually advancing from the theta peak as the animal crosses the place field, a phenomenon known as phase precession (O’Keefe and Recce, 1993) (Figure 2A). The discovery of these two neurophysiological correlates raised a debate on whether the brain uses rate and/or temporal codes for space, and whether these codes would be mechanistically related or independent (Cei et al., 2014; Harris et al., 2002; Huxter et al., 2003; Mehta et al., 2002). In the present work, we revisited this debate by investigating the evolution of theta-phase coupling of spikes along space.

Place cells code for space by precisely timing their firing at specific phases of the theta cycle (Jensen and Lisman, 2000; O’Keefe and Recce, 1993); therefore, the study of temporal coding intrinsically relates to the study of theta-phase coupling. However, up to now researchers have overlooked the relation between place cell spiking and theta phase at locations preceding the entrance of the animal in the cell’s place field, probably due to the low firing rate at these locations (O’Keefe and Burgess, 2005; O’Keefe and Dostrovsky, 1971) (Figure 2C). Here we performed a simple, yet novel space normalization (Figure 2D) that allowed us to study the coupling of place cells to theta at positions of low firing. Surprisingly, we found that place cells align to theta phase much before they increase their firing rate, that is, much before the animal enters the rate-based place field (Figures 2E and 3B). In contrast, we found that theta-phase decoupling and firing rate decrease happen simultaneously as the animal leaves the place field. These novel analyses show that hippocampal place cells display strong asymmetry between rate and temporal codes for space, and also within temporal coding itself with respect to the place field center.

Place cells represent the current position of the animal when firing maximally at the theta trough, while low-rate spikes at the first and second halves of the theta cycle (that is, before and after the trough) code for past and upcoming positions, respectively (Skaggs et al., 1996; Dragoi and Buzsáki, 2006; Buzsáki and Moser, 2013; Sanders et al., 2015). Here we found that the coupling of place cells’ spike timing to the second half of theta anticipates the entrance of the animal into the (rate-based) place field; on the other hand, place cells’ spikes rapidly decouple from the first half of the theta cycle while the animal leaves the place field. Thus, our results indicate that the representation of future positions by temporal coding reaches farther distances than the representation of past ones. This suggests that spatial navigation requires information of the immediate past and farther representation of future goals. From a translational and speculative point of view, this result may also indicate that our brain processes more information about where we are going than from where we came.

Despite considerable progress, the exact mechanism underlying phase precession remains unknown (O’Keefe and Burgess, 2005; Maurer and McNaughton, 2007; Burgess and O’Keefe 2011). Here we provide new evidence dissociating changes in firing rate and theta-phase coupling. Although our results do not directly address phase precession mechanisms, they provide insights into previously proposed models. Influential theories of phase precession include dual oscillatory interference models (O’Keefe and Recce, 1993), ramp depolarization (Mehta et al., 2002), and models based on internal network dynamics (Maurer and McNaughton, 2007). Dual oscillatory models posit that phase precession is due to the interference of two oscillators of slightly different theta frequencies: one at the LFP theta frequency generated at the soma, and another at faster frequency generated at the dendrites (O’Keefe and Burgess, 2005; Burgess and O’Keefe 2011). The oscillatory interference gives rise to membrane potential oscillations that change spike-phase probability across theta cycles (Burgess and O’Keefe 2011). However, the current models of oscillatory interference would not account for the different levels of phase locking at similar firing rate values observed here (i.e., when entering vs leaving the place field). The ramp depolarization model also assumes a theta-oscillating membrane potential but posits that phase precession is due to a monotonically increasing excitatory drive as the animal enters the place field (Mehta et al., 2002); this model was motivated by the negative skewness of rate-based place fields originally reported in Mehta et al. (2000). In principle, such a mechanism could account for high spike-phase locking preceding entrance into the rate-based place field, assuming that low membrane depolarizations may be just enough to induce spiking locked to theta without major increases in firing rate. However, firing rate asymmetry was rather low in the present dataset (Figures S3 and S4; see also Lee et al. (2004) and Park and Lee (2016) for similar skewness estimates); but perhaps more importantly, we obtained the same results when restricting our analysis to place cells exhibiting positive skewness (Figure S3), which therefore argues against a ramp depolarization model. Finally, a different class of models shows that phase precession can arise from network interactions rather than intrinsic cell properties (Maurer and McNaughton, 2007; Burgess and O’Keefe 2011); for example, a computational model has suggested that phase precession is due to asymmetric synaptic connections among place cells (Tsodyks et al., 1996). Whether network models can account for our results remains to be tested.

In summary, temporal coding by place cells has large asymmetry around the center of the rate-based place field. Our findings further revealed a three-stage dynamics for spatial coding through spike timing, in which place cells sequentially undergo phase coupling, phase precession and phase decoupling. While previous studies linked phase precession to changes in firing rate (Cei et al., 2014; Harris et al., 2002; Mehta et al., 2002), here we found that phase coupling and decoupling are independent from the latter, and moreover happen at constant but opposite theta phases. Finally, the asymmetry of temporal coding towards the beginning of the place field suggests that place cells code more for future than past locations. We conclude that the network mechanisms controlling spike timing and frequency differ, and that place cells display temporal coding before rate coding.

## MATERIAL AND METHODS

*Data set*. We analyzed CA1 LFPs and spikes recorded with silicon-based multi-site probes from 3 male Long-Evans rats navigating in a 250-cm long linear track. This dataset is freely available online at the Collaborative Research in Computational Neuroscience website (https://crcns.org/), and was generously contributed by Gyorgy Buzsáki’s laboratory (Mizuseki et al., 2013). Detailed descriptions of the experimental procedures have been largely documented elsewhere (Diba and Buzsáki, 2008; Mizuseki et al., 2009, 2014). Briefly, animals were implanted with a 4- or 8-shank silicon probe in the right dorsal hippocampus; shanks were 200 μm apart and had 8 recording sites (Fujisawa et al., 2008) (160 μm^2^, 1-3- MΩ impedance) separated by 20 μm. Signals were acquired on a 128-channel DataMax system at 20 kHz. Waveforms were sorted using KlustaKwik (Harris et al., 2000) and manually adjusted with Kluster (Hazan et al., 2006). LFPs were obtained by down-sampling to 1250 Hz. Recording sites were identified *a posteriori* using histological data, electrophysiological benchmarks, and stereotaxic coordinates. Animals were videorecorded at 39.06 Hz; position on the linear track was estimated using two light-emitting diodes placed on the top of the head.

*Data analysis*. All analyses were performed using built-in and custom written routines in MATLAB. For each shank and session, the reference LFP was selected as the channel with the highest percentage power in the theta range (5-12 Hz). Filtering was achieved by means of a finite impulse response filter from the EEGLAB toolbox (Delorme and Makeig, 2004). The instantaneous phase was obtained using the analytical representation of the signal based on the Hilbert transform.

*Place cells and place fields*. We analyzed 100 sessions across the three animals. On each recording session, left and right runs were considered independently (McNaughton et al., 1983; Muller et al., 1994). We binned the linear track in 5-cm bins and calculated the spatial information per spike as described in Skaggs et al. (1993). Units with more than 1 bit of spatial information and with global firing rate higher than 0.3 Hz were considered putative place cells. We then computed continuous spatial firing rates by smoothing spike counts and spatial occupancy with a Gaussian kernel function (SD, 5 cm). Place fields were defined as contiguous regions (> 20 cm) of firing rate above a threshold automatically set as half the average of the 50% highest firing rate bins (adapted from Senior et al., 2008). Place fields at the ends of the track (first and last 10 cm) were excluded from the analyses. Bimodal unidirectional place fields and bidirectional place fields (>50% field overlap between left and right runs) were considered as a single place field sample. Following these criteria, we obtained a total of 689 place cells and 1071 place fields.

*Phase coupling and normalized firing rate*. To calculate spike-phase coupling strength as a function of space, we binned theta phase and relative distance to the place field center in non-overlapping bins of 20° and 0.1 place field length, respectively. At each space bin, spike-phase coupling was defined as a distance metric of the empirical spike-phase distribution from the uniform distribution, as previously described (Hurtado et al., 2004; Tort et al., 2007). Theta-phase coupling strength (TPC) was computed using the same metric but applied to the distribution of mean spiking phases. Therefore, while spike-phase coupling measures theta coupling of grouped spikes, TPC estimates the consistency of the mean theta phase of spiking across place fields. To delimit the region of significant TPC values in Figure 3A,B, we generated a distribution of 1000 surrogate TPC curves, obtained by randomizing the mean spiking phases within space bins. The statistical threshold was set as the 99^th^ percentile of the surrogate distribution. In Figures 3 and 4, the TPC curve was corrected by subtracting the mean surrogate curve. To compute the mean normalized firing rate (FR) curve, for each place field we divided the spatial firing rate by its maximum. TPC and FR curves were smoothed using a cubic spline before assessing the positions of peak and half maximum values; 99% confidence intervals for these parameters were obtained using 1000 random subsamples of 70% of place fields (Efron, 1981).

## AUTHOR CONTRIBUTIONS

A.B.L.T designed study; B.C.S. analyzed data under supervision of A.B.L.T.; A.B.L.T. and B.C.S. wrote the manuscript.

## ACKNOWLEDGEMENTS

We thank the Buzsáki laboratory for making data publicly available at http://crcns.org/, a data-sharing website supported by NSF and NIH, USA. We thank D. Laplagne, O. Amaral, C. Rennó-Costa, A. Lockmann, R. Pavão, V. Lopes-dos-Santos, H. Belchior, and R. Scheffer-Teixeira for critical discussions and comments on the manuscript. A.B.L.T. and B.C.S. were supported by CNPq and CAPES, Brazil. The authors declare no competing financial interests.

## Supplemental Information

**Figure S1.**
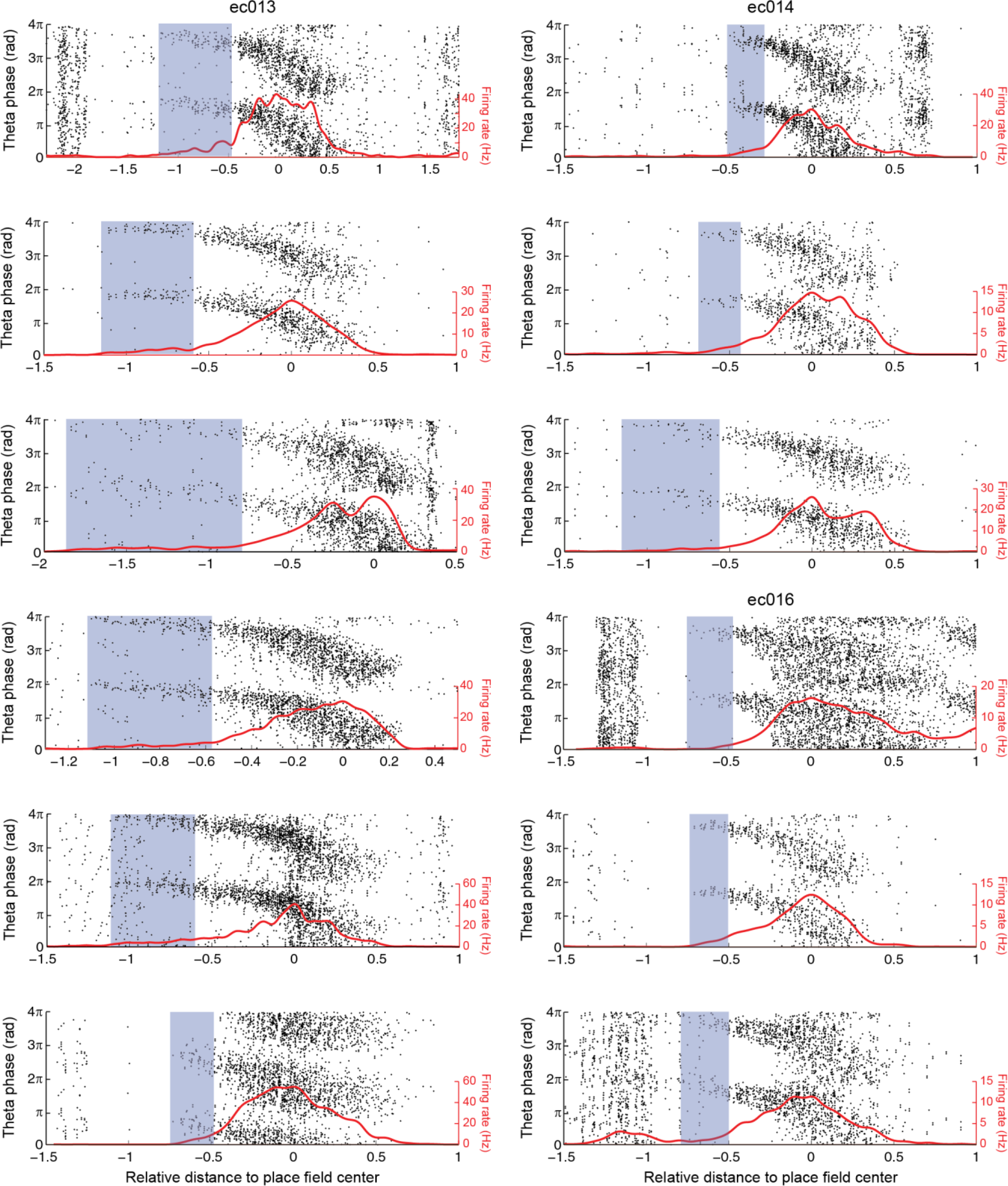
Examples of theta-phase coupling preceeding major increases in firing rate in individual place cells. Blue shaded areas highlight locations where spikes align to a preferred theta phase before major increases in firing rate. Panels on the left column were obtained for example place cells recorded from rat ec013; top and bottom three panels on the right column were obtained for example cells from rats ec014 and ec016, respectively.

**Figure S2.**
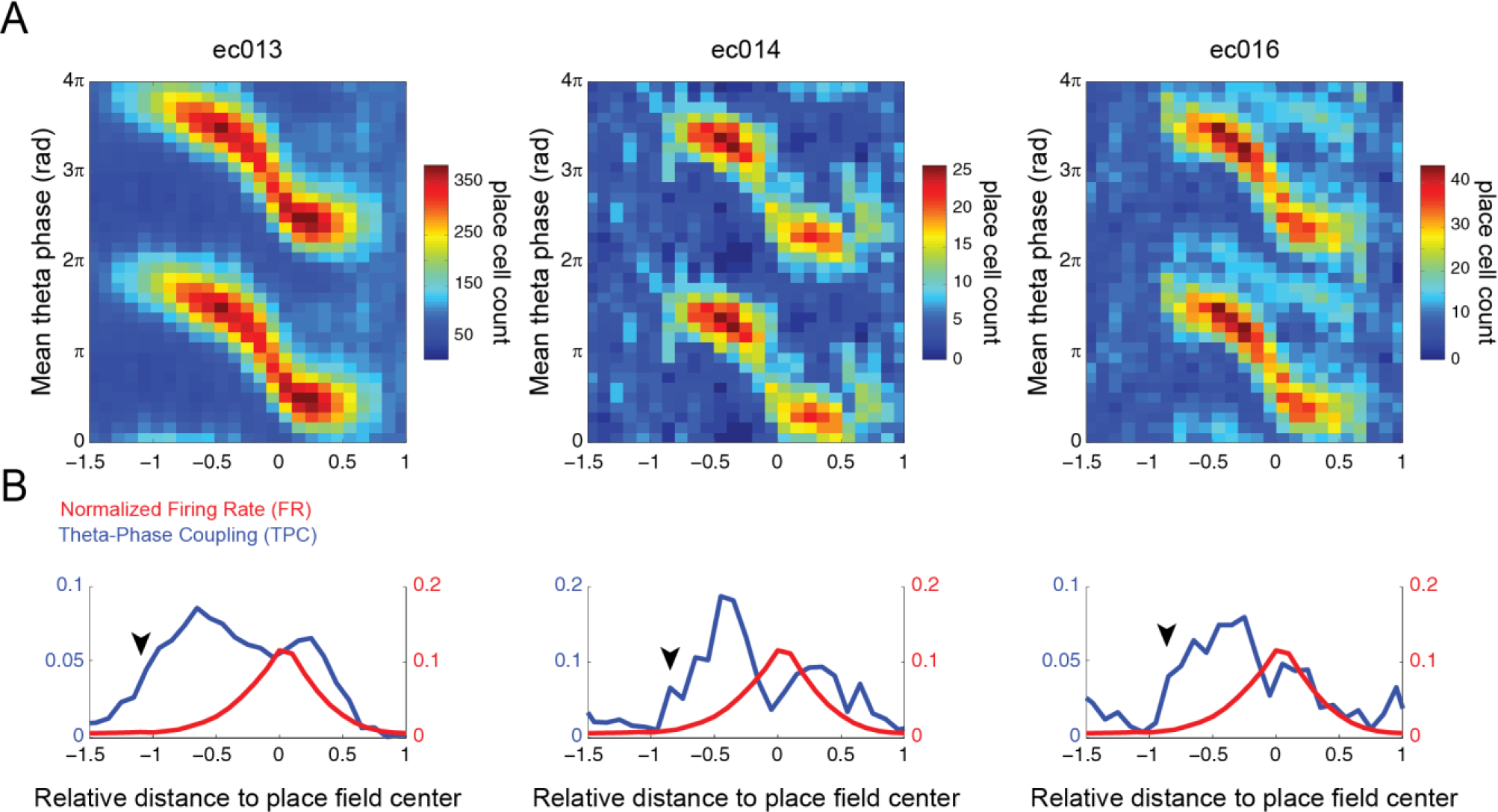
Asymmetry of temporal coding in individual animals. (A) Histograms of mean spiking phase, computed separately for each animal. At each position, individual place fields contribute with one count to the phase bin of their place cell’s mean spiking. (B) Theta-phase coupling strength (TPC, blue) and mean normalized firing rate (FR, red) as functions of space. Arrowheads point to locations where place cells couple to theta before major changes in firing rate.

**Figure S3.**
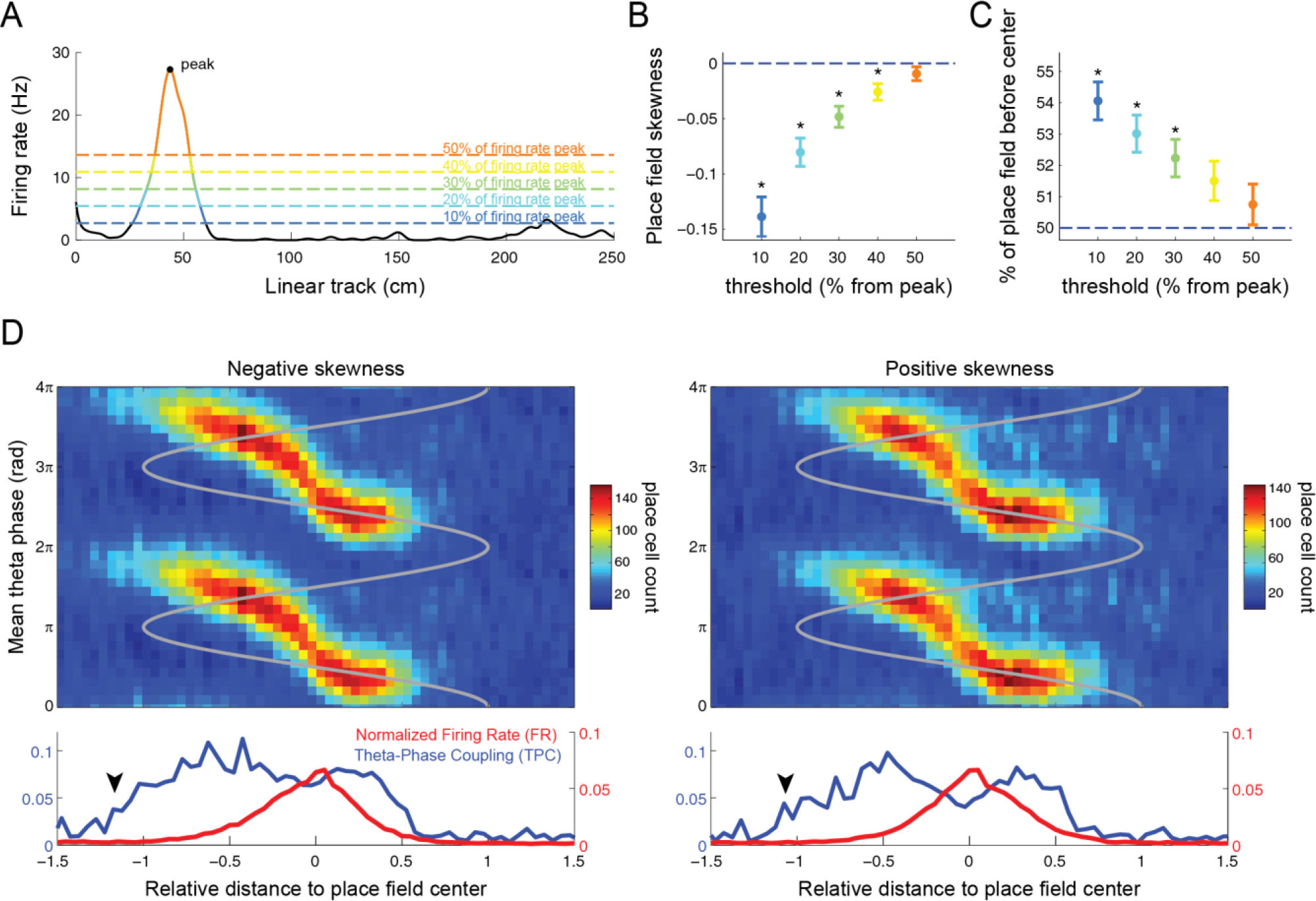
Asymmetric temporal coding irrespective of place field skewness. (A) Example place field. Dashed lines show different firing rate thresholds (at 10, 20, 30, 40 and 50% of the peak value) that were used to quantify place field asymmetry. (B) Skewness of the firing rate curve at each threshold. (C) Percentage of the place field located before the center (position of peak firing rate). In B-C, data points show the mean over individual place fields; errorbars denote standard error of the mean (*p<0.01, t-test against 0 or 50% corrected for multiple comparisons). (D) Panels show the same as in Figure S2, but computed using only place cells with negative (left) or positive (right) skewness at the 50% threshold

**Figure S4.**
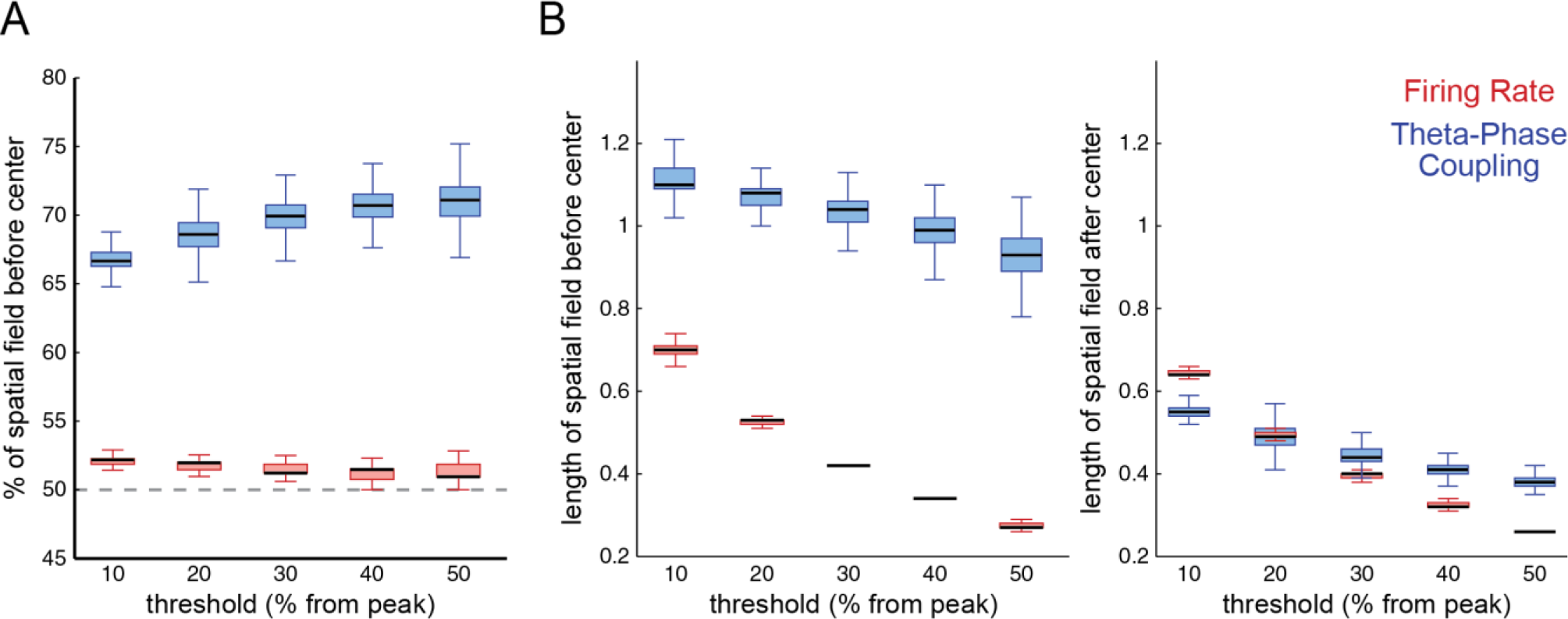
Temporal coding has greater asymmetry around the place field center than rate coding. (A) Percentage of the spatial receptive field before the place field center for TPC (blue) and FR (red) curves computed as in Figure 3F but for different thresholds (10, 20, 30, 40, and 50% of the TPC or FR peak value). Notice larger asymmetry for TPC spatial fields at all thresholds. (B) Length of the TPC and FR spatial fields before (left) and after (right) the place field center for different thresholds.

## REFERENCES

Bose, A., Booth, V., Recce, M. (2000). A temporal mechanism for generating the phase precession of hippocampal place cells. J. Comput. Neurosci. 9, 5–30.

Burgess, N., O’Keefe, J. (2011). Models of place and grid cell firing and theta rhythmicity. Curr. Opin. Neurobiol. 21, 734–744.

Buzsáki, G., Moser, E.I. (2013). Memory, navigation and theta rhythm in the hippocampal-entorhinal system. Nat. Neurosci. 16, 130–138.

Cei, A., Girardeau, G., Drieu, C., El Kanbi, K., Zugaro, M. (2014). Reversed theta sequences of hippocampal cell assemblies during backward travel. Nat. Neurosci. 17, 719–724.

Delorme, A., Makeig, S. (2004). EEGLAB: an open source toolbox for analysis of singletrial EEG dynamics including independent component analysis. J. Neurosci. Methods 134, 9–21.

Diba, K., Buzsáki, G. (2008). Hippocampal network dynamics constrain the time lag between pyramidal cells across modified environments. J. Neurosci. 28, 13448–13456.

Dragoi, G., Buzsáki, G. (2006). Temporal encoding of place sequences by hippocampal cell assemblies. Neuron 50, 145–157.

Efron, B. (1981). Nonparametric estimates of standard error: the jackknife, the bootstrap and other methods. Biometrika 68, 589–599.

Fujisawa, S., Amarasingham, A., Harrison, M.T., Buzsáki, G. (2008). Behavior-dependent short-term assembly dynamics in the medial prefrontal cortex. Nat. Neurosci. 11, 823–833.

Harris, K.D., Henze, D.A., Csicsvari, J., Hirase, H., Buzsáki, G. (2000). Accuracy of tetrode spike separation as determined by simultaneous intracellular and extracellular measurements. J. Neurophysiol. 84, 401–414.

Harris, K.D., Henze, D.A., Hirase, H., Leinekugel, X., Dragoi, G., Czurkó, A., Buzsáki, G., 2002. Spike train dynamics predicts theta-related phase precession in hippocampal pyramidal cells. Nature 417, 738–741.

Hazan, L., Zugaro, M., Buzsáki, G. (2006). Klusters, NeuroScope, NDManager: a free software suite for neurophysiological data processing and visualization. J. Neurosci. Methods 155, 207–216.

Hurtado, J.M., Rubchinsky, L.L., Sigvardt, K.A. (2004). Statistical method for detection of phase-locking episodes in neural oscillations. J. Neurophysiol. 91, 1883–1898.

Huxter, J., Burgess, N., O’Keefe, J. (2003). Independent rate and temporal coding in hippocampal pyramidal cells. Nature 425, 828–832.

Jensen, O., Lisman, J.E. (2000). Position reconstruction from an ensemble of hippocampal place cells: contribution of theta phase coding. J. Neurophysiol. 83, 2602–2609.

Lee, I., Rao, G., Knierim, J. (2004). A double dissociation between hippocampal subfields: differential time course of CA3 and CA1 place cells for processing changed environments. Neuron 5, 803–815.

Maurer, A.P., McNaughton, B.L. (2007). Network and intrinsic cellular mechanisms underlying theta phase precession of hippocampal neurons. Trends Neurosci. 7, 325–333.

McNaughton, B.L., Barnes, C.A., O’Keefe, J. (1983). The contributions of position, direction, and velocity to single unit activity in the hippocampus of freely-moving rats. Exp. Brain Res. 52, 41–49.

Mehta, M.R., Quirk, M.C., Wilson, M.A. (2000). Experience-dependent asymmetric shape of hippocampal receptive fields. Neuron 25, 707–715.

Mehta, M.R., Lee, A.K., Wilson, M.A. (2002). Role of experience and oscillations in transforming a rate code into a temporal code. Nature 417, 741–746.

Mizuseki, K., Diba, K., Pastalkova, E., Teeters, J., Sirota, A., Buzsáki, G. (2014). Neurosharing: large-scale data sets (spike, LFP) recorded from the hippocampal-entorhinal system in behaving rats. F1000Res 3.

Mizuseki, K., Sirota, A., Pastalkova, E., Buzsáki, G. (2009). Theta oscillations provide temporal windows for local circuit computation in the entorhinal-hippocampal loop. Neuron 64, 267–280.

Mizuseki, K., Sirota, A., Pastalkova, E., Diba, K., Buzsáki, G. (2013). Multiple single unit recordings from different rat hippocampal and entorhinal regions while the animals were performing multiple behavioral tasks. CRCNS Org.

Morris, R.G.M., Garrud, P., Rawlins, J.N.P., O’Keefe, J. (1982). Place navigation impaired in rats with hippocampal lesions. Nature 297, 681–683.

Muller, R.U., Bostock, E., Taube, J.S., Kubie, J.L. (1994). On the directional firing properties of hippocampal place cells. J. Neurosci. 14, 7235–7251.

O’Keefe, J., Burgess, N. (2005). Dual phase and rate coding in hippocampal place cells: theoretical significance and relationship to entorhinal grid cells. Hippocampus 15, 853–866.

O’Keefe, J., Dostrovsky, J. (1971). The hippocampus as a spatial map. Preliminary evidence from unit activity in the freely-moving rat. Brain Res. 34, 171–175.

O’Keefe, J., Recce, M.L. (1993). Phase relationship between hippocampal place units and the EEG theta rhythm. Hippocampus 3, 317–330.

Park, S.B., Lee, I. (2016). Increased variability and asymmetric expansion of the hippocampal spatial representation in a distal cue-dependent memory task. Hippocampus ahead of print.

Sanders, H., Rennó-Costa, C., Idiart, M., Lisman, J. (2015). Grid cells and place cells: an integrated view of their navigational and memory function. Trends Neurosci. 38, 763–775.

Senior, T.J., Huxter, J.R., Allen, K., O’Neill, J., Csicsvari, J. (2008). Gamma oscillatory firing reveals distinct populations of pyramidal cells in the CA1 region of the hippocampus. J. Neurosci. 28, 2274–2286.

Skaggs, W.E., McNaughton, B.L., Gothard, K.M., Markus, E.J. (1993). An information-theoretic approach to deciphering the hippocampal code. In Advances in Neural Information Processing Systems 5 (eds.Hanson, S.J., Giles, C.L. &Cowan, J.D.), pp. 1030–1037.

Skaggs, W.E., McNaughton, B.L., Wilson, M.A., Barnes, C.A. (1996). Theta phase precession in hippocampal neuronal populations and the compression of temporal sequences. Hippocampus 6, 149–172.

Tort, A.B.L., Rotstein, H.G., Dugladze, T., Gloveli, T., Kopell, N.J. (2007). On the formation of gamma-coherent cell assemblies by oriens lacunosum-moleculare interneurons in the hippocampus. Proc. Natl. Acad. Sci. 104, 13490–13495.

Tsodyks, M.V., Skaggs, W.E., Sejnowski, T.J., McNaughton, B.L. (1996). Population dynamics and theta rhythm phase precession of hippocampal place cell firing: a spiking neuron model. Hippocampus 6, 271–280.

Wilson, M.A., McNaughton, B.L. (1993). Dynamics of the hippocampal ensemble code for space. Science 261, 1055–1058.

